# The human cerebellum encodes temporally sensitive reinforcement learning signals

**DOI:** 10.1101/2025.09.06.674658

**Authors:** Juliana E. Trach, Yiran Ou, Samuel D. McDougle

**Author notes:** Correspondence: Juliana E. Trach.

## Abstract

In addition to supervised motor learning, the cerebellum also supports nonmotor forms of learning, including reinforcement learning (RL). Recent studies in animal models have identified core RL signals related to reward processing, reward prediction, and prediction errors in specific regions in cerebellar cortex. However, the constraints on these signals remain poorly understood, particularly in humans. Here, we investigated cerebellar RL signals in a computationally-driven fMRI study. Human participants performed an RL task without low-level sensorimotor contingencies. We observed robust RL signals related to reward processing and reward prediction errors in cognitive regions of the cerebellum (Crus I and II). These signals were not explained by oculomotor or physiological confounds. By manipulating the delay between choices and reward outcomes, we discovered that cerebellar RL signals are temporally sensitive: robust when feedback was delivered shortly following choices, but undetectable at supra-second feedback delays. Similar delay effects were not found in other areas implicated in reward processing, including the ventral striatum and hippocampus. Further, reward prediction error activity in the cerebellum was related to behavioral performance when feedback was delivered promptly, but not when it was delayed. Connectivity analyses revealed that during RL feedback, cognitive areas of the cerebellum coactivate with a network that includes the medial and lateral prefrontal cortex and caudate nucleus. Together, these results highlight a temporally constrained contribution of the human cerebellum to a cognitive learning task.

## Introduction

A recent paradigm shift in cerebellar neuroscience recognizes that the cerebellum contributes to both motor control and cognition [1–7]. This shift is motivated by converging lines of evidence: The cerebellum is implicated in a range of cognitive neurodevelopmental disorders [8], it is reliably activated by nonmotor tasks [9,10], damage to the region produces both motor and cognitive deficits [11–16], and it has robust bidirectional anatomical connections with nonmotor areas of the neocortex and other subcortical structures like the basal ganglia [2,17–25]. Despite this evidence, a unifying framework of cerebellar function has been elusive. What specific cognitive computations might the cerebellum carry out?

Recent work on this question has focused on cerebellar contributions to a domain that cuts across both cognitive and motor behaviors: reinforcement learning (RL) [26,27]. Studies in rodents and nonhuman primates have revealed cerebellar signals related to core components of RL, including reward processing, reward expectation, and reward prediction errors (the difference between expected and received reward) [26,28–31]. These RL signals have been observed across the cerebellar circuit, including in granule cells [30], climbing fibers [29,32,33], and Purkinje cells [31]. In particular, Crus I and Crus II regions have been highlighted as cerebellar loci for RL signals in both rodents and nonhuman primates [31,32]. Moreover, lesion and stimulation studies in animal models and humans point to a causal role for the cerebellum in reward-guided behavior, even in nonmotor tasks [12,34–36]. There is now a general consensus that one key function of the cerebellum in cognition may be to support or enhance RL [37], in conjunction with established RL circuits in the midbrain, basal ganglia, and prefrontal cortex.

Despite these discoveries, key gaps remain. First, there are practical issues: In humans, full coverage of the cerebellum with functional neuroimaging is rarely achieved or even attempted [38], and cerebellar BOLD signals are susceptible to motor and physiological confounds [39–41]. Second, many RL tasks involve sensorimotor contingencies (e.g., where specific limb movements are associated with specific rewards, or where reward feedback predictably elicits movement), making it hard to disentangle cerebellar contributions to cognitive components of RL versus sensorimotor learning [31,42,43]. Finally, in the case of incidental findings of RL-related activity in the cerebellum in humans, specific anatomical localization (other than hemispheric differences) of such signals is rarely reported and the tasks are quite variable, making such findings difficult to compare to work in model organisms. These challenges may explain heterogeneity in previously reported cerebellar activations related to reward processing [42,44,45].

There are also theoretical gaps in our understanding of cerebellar involvement in RL, primarily in the lack of known constraints on cerebellar RL computations. One candidate constraint is the cerebellum’s temporal sensitivity. In tasks like eyeblink conditioning and motor adaptation, the cerebellum is optimized for cue-outcome associations separated by short intervals (i.e., < ∼2 seconds) [46–51], and typically does not contribute to associative learning when these intervals are exceeded [52]. In contrast, areas like the basal ganglia, hippocampus, and neocortex connect events over longer timescales [53,54]. We thus hypothesize that the cerebellum may be selectively engaged in RL contexts where predictive associations unfold over short intervals.

We directly tested this hypothesis using choice behavior, eye-tracking, physiological recordings, computational modeling, and whole-brain model-based fMRI in human subjects. Our RL task avoided low-level sensorimotor associations, and we optimized cerebellar coverage and accounted for several potential confounds. By manipulating the delay between choice and reward feedback we investigated if and how the choice-outcome interval modulates cerebellar contributions to RL. Moreover, the flexibility of whole-brain fMRI allowed us to ask how RL signals in the cerebellum compare to those seen in canonical subcortical and cortical RL regions, to link neural responses to behavioral outcomes, and to ask if RL feedback selectively drives functional correlations between the cerebellum and other regions of the brain.

## Results

### Human Crus I and Crus II preferentially respond to reward feedback

Participants (N=32) performed a probabilistic reinforcement learning (RL) task while undergoing fMRI. Participants observed two stimuli on each trial and used a button box to select the stimulus (left or right) that they thought was most likely to yield a reward (**Figure 1a**). Their goal was to win points to earn a monetary bonus at the end of the task. After they made their choice, there was a delay period (short delay: 0.8s; long delay: 3s) before they received probabilistic reward feedback (nonrewarded trials: +0; rewarded trials: +1) which they could use to guide future choices. Participants saw four distinct pairs of stimuli within each run (see *Methods* and **Supplemental Figure 1**). In each pair, one stimulus was associated with a .75 probability of reward and the other with a .25 probability of reward. Importantly, the location of the stimuli was randomized across trials, such that rewards were associated with the abstract stimulus choice rather than a specific motor action or spatial location. Based on previous work, our *a priori* regions of interest (ROIs) were Crus I and Crus II in the cerebellum (following work in nonhuman primates [31] and rodents [32]), and canonical subcortical RL regions in nucleus accumbens (NAc), caudate nucleus (Cd), and hippocampus (Hpc; **Figure 1b**).

**Figure 1.**
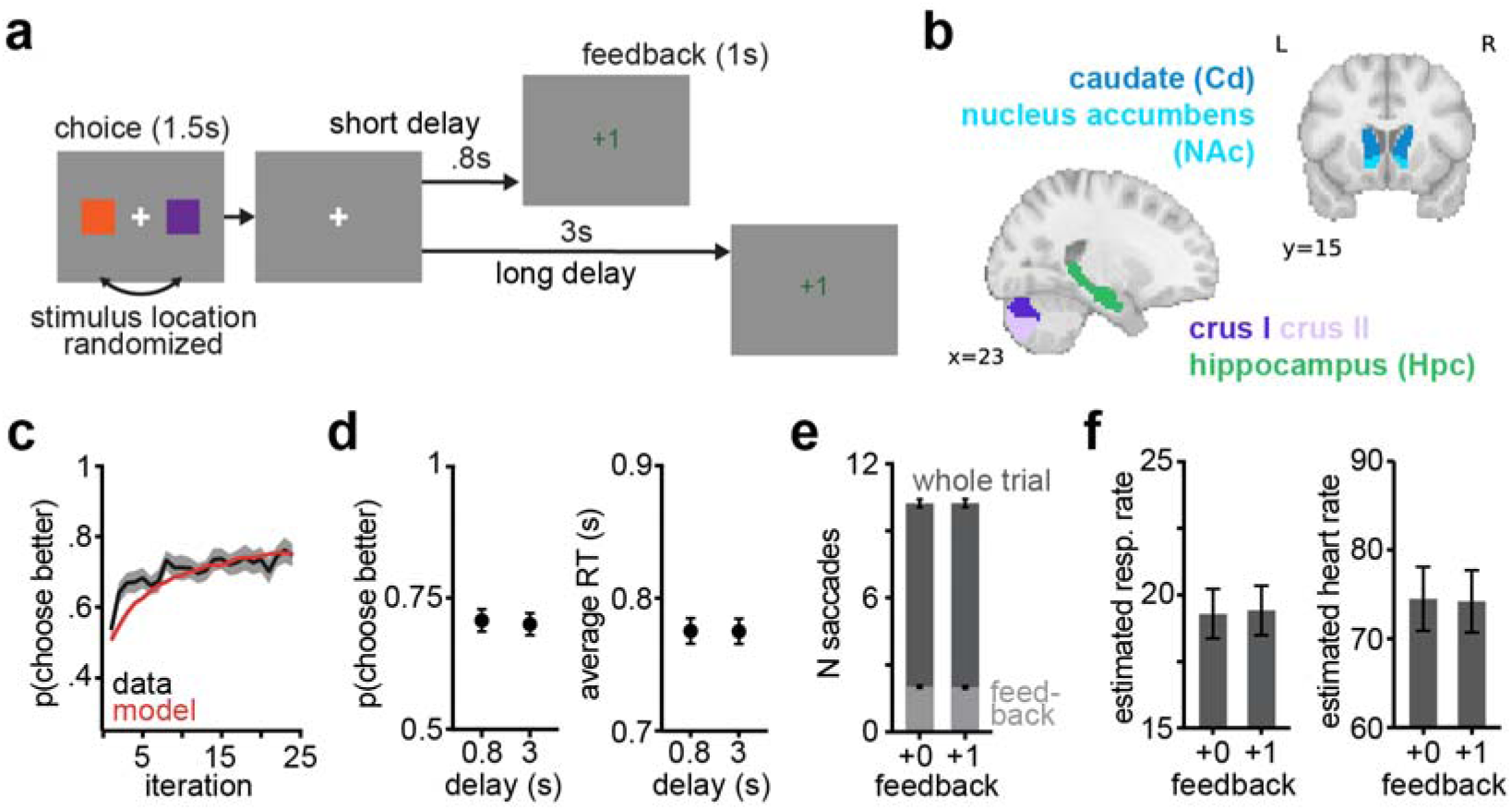
a) Task schematic. Example stimuli pictured in Supplemental Figure 1. b) Anatomical ROI masks. c) Learning curve and model fit. d) Choice and RT performance on short and long delay trials. e) Gaze on nonrewarded (+0) versus rewarded (+1) trials. Larger values depict data from whole trial and smaller values depict data during the feedback phase only. f) Respiration and pulse rates on rewarded versus nonrewarded trials.

Participants performed the RL task well, showing significant learning across trial blocks (GLM: choose correct ∼ iteration + (1|subject); iteration: *b* = 0.026, *SE* = 0.0037, *z* = 6.93, *p* < .001; **Figure 1c**). These learning data were fit with an RL model which was used for later model-based fMRI analyses (see *Methods*). At the group level, feedback delay did not significantly affect choice behavior or reaction times (p(choose higher value stimulus): *t*(31) = 0.51, 95% CI = [-0.02, 0.04], *p* = .616; RT: *t*(31) = 0.03, 95% CI = [-0.01, 0.01], *p* = .976; **Figure 1d**). Crucially, there was comparable gaze behavior (**Figure 1e**; at feedback: N saccades: *t*(28) = 0.76, 95% CI = [-0.05, 0.11], *p* = .451) and physiological measures (**Figure 1f**; rewarded versus nonrewarded trials: respiration: *t*(23) = −0.95, 95% CI = [-0.37, 0.14], *p* = .353; pulse: *t*(18) = 1.27, 95% CI = [-0.25, 0.83], *p* = .274) on rewarded and nonrewarded trials, meaning that our key neural analyses would not be significantly confounded by these factors. (This is an important control, as confounding factors like eye movements can influence cerebellum activity.)

For our neural analyses, we first used generalized linear models (GLMs) to examine activity between rewarded and nonrewarded trials. In addition to canonical reward processing regions (**Figure 2a, left panel**), we also observed significant activity in the lateral cerebellum around the Crus I/Crus II boundary that reflected greater feedback responses on rewarded versus nonrewarded trials (**Figure 2a, right panel**). The localization of these signals corresponds with previously reported effects in both rodents [32] and nonhuman primates [31].

**Figure 2.**
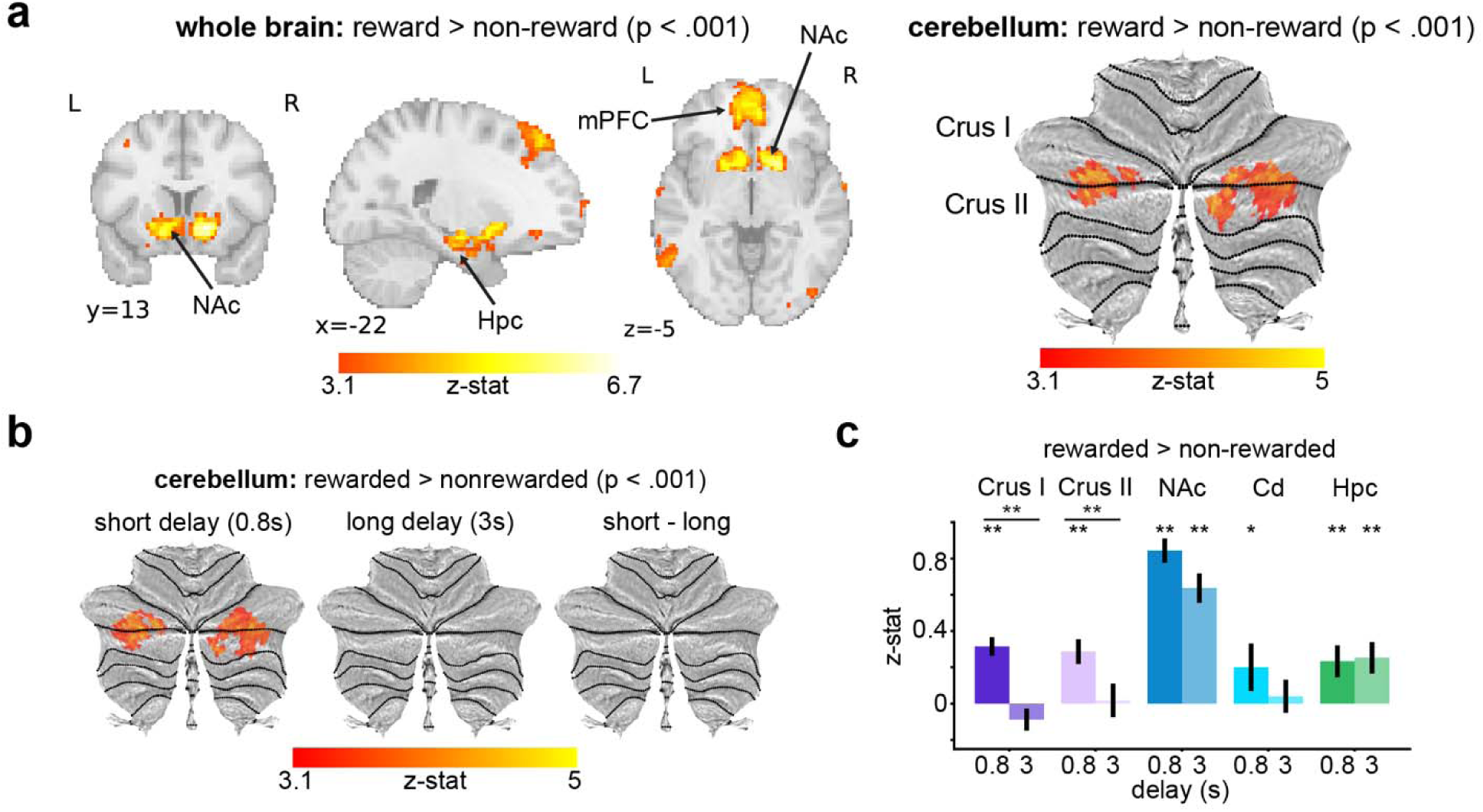
a) Contrast of activity on rewarded versus nonrewarded trials at the point of feedback in the whole brain (left panel) and in cerebellum (right panel; plotted on cerebellar flat map, see Methods). Whole brain result is cluster-corrected at p < .001. Cerebellar results are cluster-corrected in cerebellum, p < .001. b) Rewarded > nonrewarded contrast at short and long delays in cerebellum (cluster-corrected in cerebellum, p < .001). c) Contrast of activity for rewarded > non-rewarded trials in cerebellar (Crus I and II), striatal (Nucleus Accumbens, NAc; Caudate Nucleus, Cd), and hippocampal (Hpc) ROIs. ** p < .001; * p < .05 Cerebellar reward signals are temporally constrained

We next tested whether these cerebellar signals were affected by feedback delay. We hypothesized that the cerebellum would be specifically responsive to feedback on short-delay (0.8s) versus long-delay (3s) trials, echoing temporal constraints on cerebellar processing in motor learning tasks [49,51,55]. We thus examined responses to rewards at short- and long-delay intervals separately. In line with our hypothesis, we found significant cerebellar responses to rewards on short-delay, but not long-delay, trials (**Figure 2b**; see **Supplemental Figure 2** for whole brain results). Activity was primarily localized to Crus I and Crus II. The null findings on long delay trials held even with a relaxed statistical threshold (cluster-corrected in cerebellum, p < .05), indicating that the lack of significant reward responses at long delays was not simply a result of conservative thresholding.

We additionally asked whether this temporal sensitivity was specific to the cerebellum, or was also reflected in other reward processing regions (NAc, Cd, and Hpc; [54]). We extracted /J-values for the rewarded > nonrewarded contrast (**Figure 2c**) from these ROIs at short and long delays. Corroborating the whole cerebellum results, the *a priori* anatomical cerebellar ROIs only exhibited significant positive responses at short delays (nonparametric bootstrap test: short delays: Crus I: *M* = 0.32, 95% CI = [0.19, 0.45], *p* < .001; Crus II: *M* = 0.29, 95% CI = [0.16, 0.42], *p* < .001; long delays: Crus I: *M* = −0.09, 95% CI = [-0.26, 0.10], *p* = .346; Crus II: *M* = 0.02, 95% CI = [-0.15, 0.19], *p* = .844). In striatum, the caudate showed a similar effect as the cerebellum (nonparametric bootstrap test: short delays: *M* = 0.20, 95% CI = [0.03, 0.36], *p* =.020; long delays: *M* = 0.04, 95% CI = [-0.11, 0.20], *p* = .648). However, the ventral striatum ROI (NAc) and hippocampus (Hpc) had significant responses at both short and long feedback delays (nonparametric bootstrap test: short delays: NAc: *M* = 0.84, 95% CI = [0.60, 1.08], *p* < .001; Hpc: *M* = 0.23, 95% CI = [0.14, 0.33], *p* < .001; long delays: NAc: *M* = 0.64, 95% CI = [0.45, 0.80], *p* < .001; Hpc: *M* = 0.25, 95% CI = [0.13, 0.36], *p* < .001). Only the cerebellar ROIs exhibited *significantly* stronger responses at short versus long feedback delays (short > long: Crus I: *M* = 0.30, 95% CI = [0.18, 0.44], *p* < .001; Crus II: *M* = 0.20, 95% CI = [0.07, 0.33], *p* = .002; Cd: *M* = 0.12, 95% CI = [-0.01, 0.25], *p* = .072; NAc: *M* = 0.15, 95% CI = [-0.01, 0.31], *p* = .072; Hpc: *M* = −0.01, 95% CI = [-0.11, 0.08], *p* = .826). Further, the difference in response to short versus long delay feedback was significantly larger in Crus I than in Cd or Hpc (Crus I: *M* = 0.40; NAc: *M* = 0.21; Cd: *M* = 0.16; Hpc: *M* = −0.02; Crus I vs NAc: 95% CI = [-0.07, 0.44], *p* = .142; Crus I vs Cd: 95% CI = [0.07, 0.41], *p* = .006; Crus I vs Hpc: 95% CI = [0.15,0.68], *p* = 0.002) and in Crus II (*M* = 0.27) than in Hpc (Crus II vs NAc: 95% CI = [-0.2,0.31], *p* = .614; Crus II vs Cd: 95% CI = [-0.1, 0.31], *p* = .298; Crus II vs Hpc: 95% CI = [0.05,0.53], *p* = 0.016). Taken together, these results provide evidence for temporally constrained reward processing in the human cerebellum during RL and also show that the delayed feedback did not globally blunt reward signals.

### Model-based analyses reveal time-sensitive RPE computations in human cerebellum

We next tested whether the human cerebellum tracked the core teaching signal in RL: reward prediction error (RPE). We leveraged computational modeling to examine RPEs in the cerebellum (**Figure 3a**). To do this, we fit RL models to each participant’s behavior to obtain learning and choice parameters for each subject (see *Methods*; **Figure 1c**). We then used these subject-specific parameters to simulate trial-by-trial RPEs based on a participant’s unique sequence of trials and choices. The time course of these computationally inferred RPEs serves as a prediction about activity in regions of the brain that may encode RPE signals. We restricted our main analyses to RPEs on rewarded trials to avoid the strong collinearity with valence signals (see *Methods;* see **Supplemental Figure 3** for RPE results across rewarded and nonrewarded trials).

**Figure 3.**
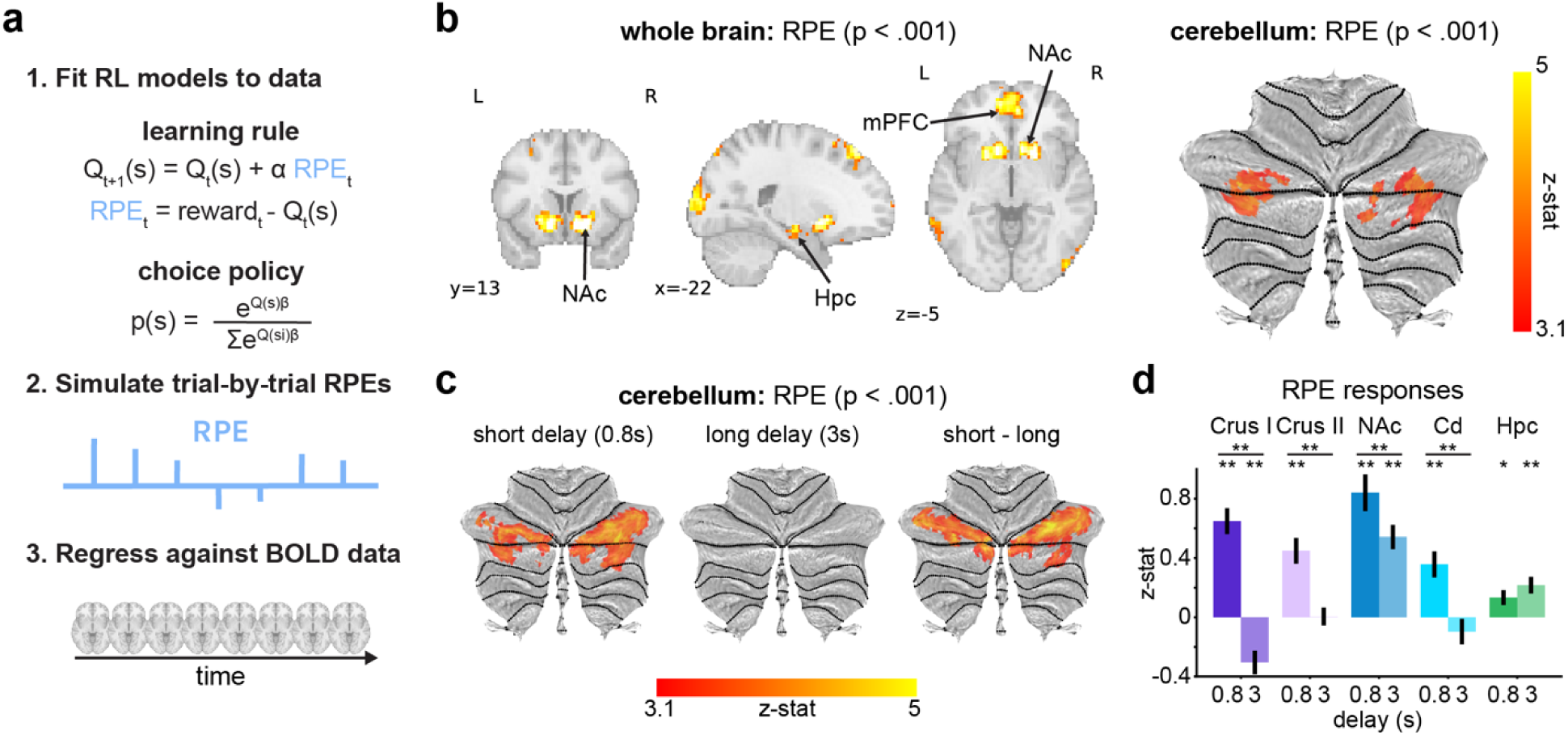
a) Schematic of model-based fMRI approach. b) Reward Prediction Error (RPE) responses (whole brain cluster-corrected, p < .001). and cerebellum (cluster-corrected in cerebellum, p < .001). c) RPE response on rewarded trials at short and long feedback delays (cluster-corrected in cerebellum, p < .001). d) RPE responses at short and long delays in cerebellar and other subcortical ROIs. **p < .001; *p < .05

We found robust RPE responses at the boundary of Crus I and Crus II in the cerebellum, in addition to canonical RPE signals in the ventral striatum (**Figure 3b**). Like reward-related activation, RPE responses were temporally constrained, and only detectable on short-delay trials (**Figure 3c** and **Supplemental Figure 3**). Crus I and II, as well as caudate, showed significant RPE responses at short, but not long delays (**Figure 3d**; nonparametric bootstrap test: short delays: Crus I: *M* = 0.65, 95% CI = [0.47, 0.81], *p* < .001; Crus II: *M* = 0.45, 95% CI = [0.28, 0.63], *p* < .001; Cd: *M* = 0.36, 95% CI = [0.20, 0.52], *p* < .001; long delays: Crus I: *M* = −0.30, 95% CI = [-0.46,-0.16], *p* < .001; Crus II: *M* = 0.01, 95% CI = [-0.10, 0.12], *p* = .886; Cd: *M* = −0.10, 95% CI = [-0.27, 0.07], *p* = .268), while nucleus accumbens and hippocampus displayed robust RPE responses at both short and long delays (short delays: NAc: *M* = 0.84, 95% CI = [0.60, 1.08], *p* < .001; Hpc: *M* = 0.13, 95% CI = [0.04, 0.23], *p* = .002; long delays: NAc: *M* = 0.54, 95% CI = [0.39, 0.70], *p* < .001; Hpc: *M* = 0.22, 95% CI = [0.11, 0.33], *p* < .001; although stronger at short delays in NAc: short > long: *M* = 0.44, 95% CI = [0.23, 0.65], *p* < .001; [54]). The difference in RPE encoding for short versus long delay feedback was significantly larger in Crus I than all other ROIs (Crus I: *M* = 0.95; Crus II: *M* = 0.44; NAc: *M* = 0.3; Cd: *M* = 0.45; Hpc: *M* = −0.08; Crus I vs NAc: 95% CI = [0.4, 0.92], *p* < .001; Crus I vs Cd: 95% CI = [0.26, 0.75], *p* < .001; Crus I vs Hpc: 95% CI = [0.78, 1.28], *p* < .001) and for Crus II versus Hpc (Crus I vs NAc: 95% CI = [-0.13,0.43], *p* = .316; Crus I vs Cd: 95% CI = [-0.28, 0.26], *p* = 0.99; Crus I vs Hpc: 95% CI = [0.29, 0.77], *p* < .001). Thus, we found that the human cerebellum encodes prediction errors that are used for learning in a nonmotor learning task, and that these teaching signals may show similar temporal constraints as those observed in sensorimotor learning.

RPEs can be signed (i.e., sensitive to valence) or unsigned (i.e., only sensitive to outcome surprise, but not valence). Thus, we used the same approach as above to examine neural correlates of surprise (|RPE|). This analysis, however, did not reveal any significant clusters of activity in the cerebellum, even at a relaxed statistical threshold, suggesting that human cerebellar RPE activity is dominated by a reward-sensitive signal (see *Discussion*).

### Linking cerebellar RPE activity to behavior

An additional question we asked is whether the apparent cerebellar involvement in RL at short (but not long) feedback delays covaries with behavioral performance in the task. We note that we did not see general performance differences between delay conditions at the group level: On average, participants showed comparable learning and RTs for stimulus pairs associated with short versus long feedback delays (p(choose higher value): *t*(31) = 0.51, 95% CI = [-0.02, 0.04], *p* = .616; RT: *t*(31) = 0.03, 95% CI = [-0.01, 0.01], *p* = .976; **Figure 1d**). However, we did observe individual differences in learning efficacy.

To investigate brain-behavior correlations, we compared RPE signals for short versus long delay trials in a whole-brain GLM, and included RL performance in the short delay condition as a participant-wise covariate in the model. In other words, this analysis tests for voxels in the brain where stronger temporally sensitive RPE responses covary with how well a participant performed in the short delay condition. Strikingly, this whole-brain analysis revealed significant brain-behavior correlations only within the cerebellum: The analysis revealed a single significant cluster of activity on the Crus I/II boundary (**Figure 4**) (albeit at a slightly relaxed statistical threshold for the whole-brain case). As a control, we also repeated this analysis using performance on the long delay condition, and the difference between performance on the short versus long delay conditions, as covariates, and observed no significant effects. Although this analysis is correlational, it suggests that better reinforcement learning (in the short delay condition only) was related to stronger cerebellar RPEs, thus linking our cerebellar results to behavioral outcomes.

**Figure 4.**
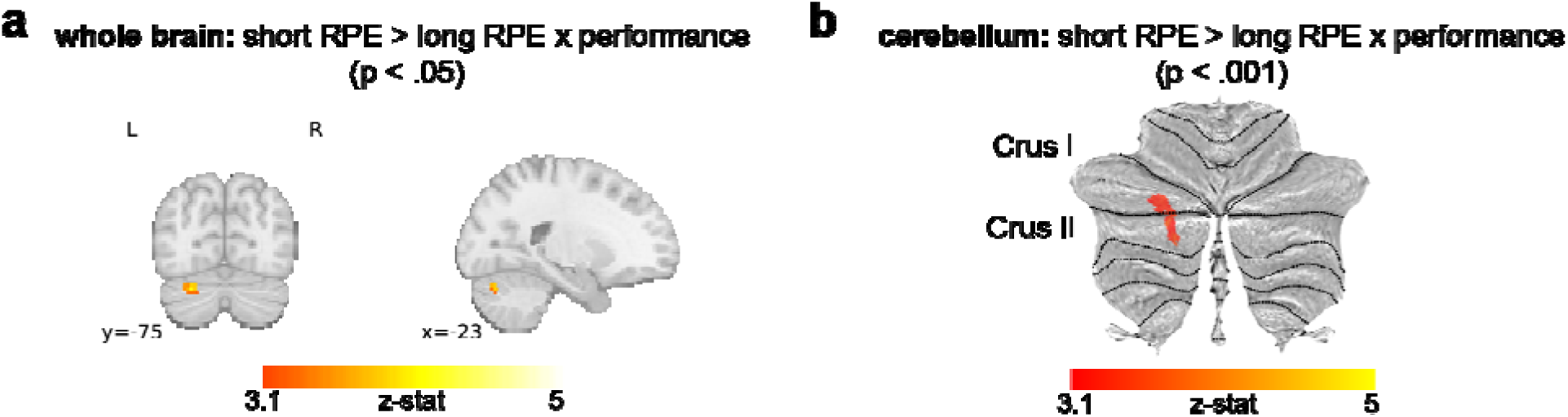
a) Whole-brain analysis showing voxels where short RPE > long RPE contrast covaries with performance on short delay trials. (cluster-corrected in whole brain, p < .05). b) Same analysis restricted to cerebellum (cluster-corrected in cerebellum, p < .001).

### Increased connectivity between cerebellar ROIs and cerebrum during feedback

The cerebellum has bidirectional connections to much of the cerebral RL circuitry, including the basal ganglia and prefrontal cortex [17,56]. Do cerebellar responses to RL feedback reflect connectivity with a wider cerebellar-striatal-frontal functional circuit? To address this question, we asked which regions of the cerebrum might be functionally coupled with the cerebellum during RL, specifically during feedback processing. We conducted exploratory psycho-physiological interaction analyses (PPI; [57]; **Figure 5a**) seeded in our two *a priori* anatomical cerebellar ROIs (Crus I and Crus II) to measure if and how functional correlations between the cerebellum and the rest of the brain increase during RL feedback above and beyond other task phases.

**Figure 5.**
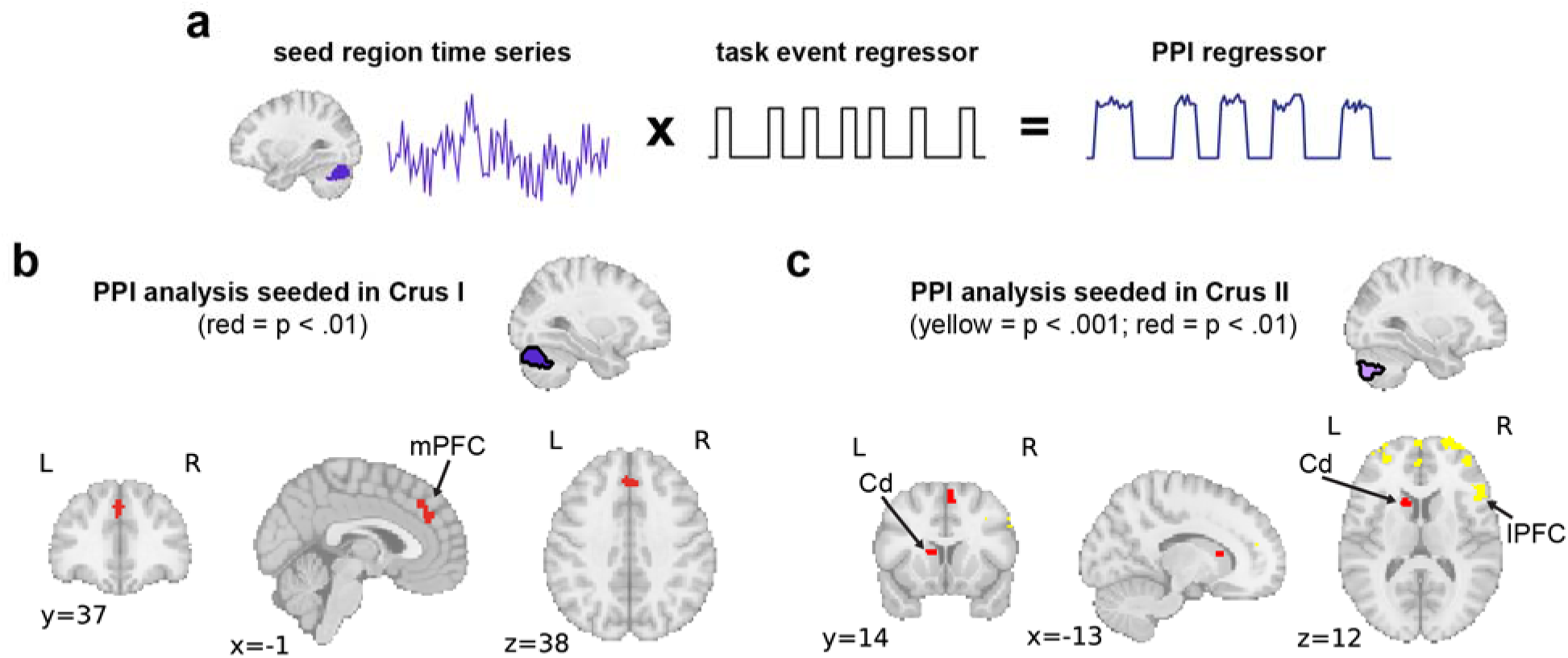
a) Illustration of PPI analysis. b) PPI analysis results seeded in Crus I (red regions cluster-corrected at p < .01). c) PPI analysis results seeded in Crus II (red cluster-corrected at p < .01, yellow at p < .001).

We observed significant activity in the medial prefrontal cortex reflecting robust feedback-sensitive functional connectivity with cognitive regions of the cerebellum during RL (**Figure 5b & c**). Crus II also showed RL feedback-related functional correlations with lateral PFC regions. Further, at a somewhat relaxed statistical threshold, we saw significant connectivity between Crus II and the caudate nucleus (**Figure 5c**), the subcortical region that showed similar temporal sensitivity for RPE responses. These results suggest that nonmotor reward feedback increases functional coupling between the cerebellum and core corticostriatal circuits that support RL.

## Discussion

Here, we present novel evidence of temporally constrained cerebellar involvement in RL in humans. We found that nonmotor regions of the human cerebellum (Crus I and II; i.e., the “cognitive” cerebellum) respond to rewards and encode trial-by-trial reward prediction errors, building on and extending recent work in rodents and non-human primates [29–33]. Consistent with known temporal constraints on cerebellar computation in the sensorimotor domain [46,48,52], we found that cerebellar RL responses are sensitive to feedback timing – delaying feedback by several seconds abolished cerebellar RPE responses, while delayed feedback signals were preserved in multiple other regions [54]. Further, the degree of RPE encoding on short versus long delay trials covaried with behavioral performance on short delay trials, suggesting that cerebellar RPEs might contribute to the efficacy of RL processes. Functional connectivity analyses revealed communication between the cerebellum and canonical corticostriatal circuitry during RL feedback, further emphasizing the cerebellum’s role as a node in the RL network [19,34,56,58]. Specifically, cerebellar RL feedback responses covaried with prefrontal cortical areas and the caudate nucleus, with the latter echoing the temporal sensitivity of cerebellar responses. Overall, these results implicate the cerebellum in a key domain of prediction error-based learning beyond supervised sensorimotor learning, and further highlight the importance of examining cerebellar responses in cognitive neuroscience [59].

We note that our results are not the first to report prediction error signals in the human cerebellum [60,61] or to suggest that the basal ganglia and cerebellum work collaboratively [58]. This previous work, however, did not necessarily isolate the signals we see here: First, in previous work in both animal models and humans, rewards are typically attached to specific motor actions, making it unclear whether cerebellar responses are driven by reward learning versus sensorimotor learning [30,62]. Similarly, in other fMRI studies reporting cerebellar prediction error responses, administered reward or punishment feedback had significant sensorimotor components, including shocking of the skin for aversive RL tasks and the ingestion of juice for reward learning [61,63]. These forms of feedback are likely to drive sensorimotor learning processes in the cerebellum (e.g., akin to eyeblink conditioning), and thus complicate interpretations of cerebellar prediction error responses. This confound may explain why previous studies showing putative RL-related responses in the cerebellum observed activity in cerebellar regions associated more with movement versus the cognitive regions we observed here.

The fact that the cerebellar RL signals we identified were temporally sensitive could also explain why similar signals have not been reported consistently in the literature. fMRI studies are generally designed to have long delays between stimuli to accommodate the slow hemodynamic response or supra-second TR measurements, and these design choices could blunt observed cerebellar involvement in learning. Additionally, much human fMRI work typically does not report what exact regions of the cerebellum (beyond hemispheric differences) are involved.

Functional mapping of the cerebellar cortex has revealed a mosaic of functional regions within the cerebellum mirroring the complexity of the cerebral cortex, and these regions respond differently to a wide range of cognitive tasks [64–66]. With these recent advances, specific localization of cerebellar signals becomes increasingly important. Notably, precision mapping research has characterized multiple distinct functional regions within Crus I and II that are involved in different types of cognitive tasks, and interconnected with distinct cortical and striatal networks [65–67]. In the context of our results, the regions of the cerebellum where we see robust RL signals (**Figures 2** and **3**) correspond to functional regions involved in social-linguistic processing and, more relevant to our task, demand regions that support active maintenance of information and working memory computations [64,65,68]. These regions also exhibit resting state connectivity with default mode areas, and both frontoparietal and corticostriatal networks [67–70].

Relatedly, our targeted approach allowed us to draw from the animal literature to make specific *a priori* predictions about localization. We found a strong correspondence between RL-sensitive regions reported in work with rodents and nonhuman primates and what we see in our human participants, despite key differences in experimental design. While model organisms afford exceptional access to recording input and output neurons in the cerebellum, cerebellar BOLD signals primarily measure *input* to the region and potentially local computations [41,71,72]. Despite this limitation, one advantage of testing human participants is that they can engage in rapid RL over a matter of minutes, respond to verbal instructions, and tolerate higher task variability across trials (e.g., multiple feedback delays), whereas model organisms require longer term training, which itself could change cerebellar responses [31]. We think this work highlights the preservation of cerebellar function across model organisms and emphasizes the importance of crosstalk between human and animal researchers. This idea is particularly relevant as cerebellar research becomes focused on “higher-order” cognitive functions [2,20,37], and as cognitive neuroscientists increasingly consider the cerebellum in their research.

One highlight of this work is that we observed consistent evidence of temporal constraints on cerebellar RL feedback processing and this temporal sensitivity was related to learning performance – cerebellar encoding of reward and RPE was essentially abolished when feedback was delayed. The idea that the cerebellum is primarily involved in sub-second coordination, associative learning, and timing is well-established in the motor domain [46,48,51,52,73,74]. This preference makes sense for a region of the brain thought to be originally adapted for short-timescale sensorimotor control and prediction [75]. At a mechanistic level, properties of granule cell firing and short-term plasticity at mossy fiber-granule cell synapses have been proposed to support sub-second timing computations characteristic of the cerebellum [73,76–78]. These mechanisms can induce timing-specific activity patterns that are then used by Purkinje cells (the main output neurons of the cerebellum) to refine actions or inform cognitive computations. There are likely biological limits that constrain what time intervals can be precisely learned by these mechanisms [79], providing a mechanistic account for why the cerebellum might be specifically attuned to encode short-timescale feedback signals.

Our connectivity analyses implicate communication between cognitive lobes of the cerebellum and neocortical (both medial and lateral prefrontal regions) and subcortical (caudate nucleus) regions during RL feedback processing. Although fMRI primarily measures cerebellar input in the BOLD signal, this communication likely involves bidirectional connections between the cerebellum and frontal and subcortical RL regions [19,25,56,58,66,69]. In particular, there is structural evidence for bidirectional connections between the cerebellum and striatum [17,19,58], the core of the reward system in the brain. A recent fMRI paper [60] focused on these connections demonstrated how cerebellar-striatal connectivity might support reward-based motor learning; in contrast to our work, however, they found that cerebellar activity primarily tracked surprise (i.e., unsigned prediction error magnitude), rather than signed RPE, as we see. Indeed, we did not find significant activation in the cerebellum for tracking surprise in our model-based analyses. This discrepancy is likely due to significant task differences, where cerebellar signals in [60] were associated with successful versus unsuccessful outcomes of a *motor* task, not a nonmotor learning task as in our study. Further, our connectivity results highlight dorsal rather than ventral regions of the striatum targeted in [60].

Overall, this work provides evidence of temporally constrained, behaviorally relevant RL signals in the human cerebellum. RPE and reward-related signals were prominent in Crus I and II – the cognitive zones of the cerebellum – providing high correspondence with recent work in animal models. Connectivity results demonstrated functional correlations between Crus I and II and regions of the medial and lateral frontal cortex, and caudate nucleus, during RL. This work extends beyond the well-studied role of the cerebellum in prediction-error based supervised motor learning [27,79,81–83], and contributes to growing evidence that the human cerebellum supports prediction-error based learning in nonmotor domains.

## Methods

### Participants

Thirty-five individuals participated in the study (Mean age = 22.9 [18–33]; N female = 24). We planned on excluding participants that were not engaging in the task (not responding to >25% of trials or only responding with one response >90% of trials) or that produced too much head motion during the scan (maximum motion over 2.5mm). No participants met our behavioral exclusion threshold. Two participants were excluded for excessive motion and an additional subject was excluded for technical issues with the data. Our final sample was 32 participants (Mean age = 22.8 [18–33]; N female = 24). The task protocol was approved by the Yale Institutional Review Board. Participants were compensated $30/hr. They were also told that they could receive up to $10 bonus payment for their performance in the task to connect the reward feedback during the task to actual reward. We gave all participants the full bonus payment.

### Experimental session

The experimental session was 2hrs long. Participants arrived and were consented and screened for metal. In the scanner, they completed three runs of a probabilistic reinforcement learning (RL) task and three runs of a statistical learning (SL) task (not discussed here). The RL task was always performed before the SL task. Each functional run began with an eye-tracker calibration phase (Eyelink 1000 Plus, Long-range monocular). After the functional runs, we collected a T1 weighted anatomical image before bringing them out of the scanner. Finally, the session concluded with a test phase for the SL task and a debriefing questionnaire. Task and protocol details are explained below.

### Reinforcement learning task

Participants performed three runs of a probabilistic reinforcement learning task in the scanner. During each run, participants saw four pairs of images (8 unique images per run) across trials (Supplemental Figure 1). On each trial, they saw one pair of images side-by-side and used an MR-safe, two-button button box with their right hand to select one of the two images (index finger to select left image, middle finger to select right image). They received probabilistic reward feedback (reward: +1 or nonreward: +0) on their response after a delay (Figure 1a). One image in each pair was associated with a .25 reward probability and the other with a .75 reward probability throughout a run of trials. Participants were instructed to select the stimulus on each trial that they thought was most likely to give them points. They were informed that their point total would be translated into a monetary bonus at the end of the session (up to $10) to motivate them to perform well and ensure that the point-feedback was rewarding. Crucially, the location of each image was counterbalanced across trials so that reward information was associated with the image – the abstract choice – and not with a specific location or motor action.

We manipulated the delay between stimulus offset and the presentation of the reward feedback in order to test the hypothesis that the cerebellum would be particularly involved in processing short-latency feedback signals. Participants viewed the pairs of stimuli for 1.5s on each trial, during which time they made their response. Once they made their response, the fixation cross changed color to indicate that their choice had been registered and the stimuli remained on the screen for the remainder of the 1.5s. The stimuli then disappeared and there was a delay before they were presented with reward feedback for 1s. To test the hypothesis that the cerebellum would be sensitive to short-latency feedback, we assigned two pairs in each run to a short feedback delay where the feedback appeared .8s after the stimulus disappeared and the other two pairs to a long feedback delay where feedback appeared after 3s. If they did not respond quickly enough, they received the feedback “Please respond faster.” There was a jittered inter-trial interval (ITI; 1-3s) after feedback presentation.

Prior to the three runs of the task, participants executed a short practice block with two pairs of stimuli to orient them to the task. There was no variation in the delay between responses and feedback in the practice block. During the main runs, participants performed 96 trials during a run (24 presentations per pair randomly interleaved). New stimuli were used in each run to prevent participants from using learned values from previous runs.

### Behavioral and physiological analysis

We used participant choices and RTs to characterize their behavior. Trials where participants did not respond were excluded from analysis. Learning was operationalized as the probability of selecting the stimulus associated with the higher reward probability across trials. To quantify learning we implemented a generalized linear model that used iteration (how many times the participant had viewed a specific pair of stimuli) to predict whether or not they chose the higher value stimulus (1 = chose higher value stimulus; 0 = chose lower value stimulus).

Overall learning was measured as the probability of selecting the higher value stimulus across all trials. We used t-tests to compare overall learning and RTs between short and long delay trials and to compare gaze and biophysiological measures on rewarded versus nonrewarded trials.

We additionally collected eyetracking data during the entirety of the scan using a long-range Eyelink 1000 plus. We did not obtain eyetracking data from two participants who used MR-safe glasses during the scan which interfered with tracking and an additional participant due to equipment malfunction. We began each run with a calibration phase. We were primarily interested in differences in gaze behavior on rewarded versus nonrewarded trials that might impact our results. To examine this possibility, we summarized gaze behavior within each trial and at feedback by counting the number of saccades in these time intervals and used a t-test to compare between rewarded and nonrewarded trials.

Finally, we collected pulse and respiration data from each participant using Siemens peripheral devices during scanning (samples every 2.5ms). We used a pulse-oximeter positioned on the index finger of the participant’s left hand to collect pulse data and a respiration belt positioned around the participant’s abdomen to track respiration. A portion of the data was lost due to signal drop out during the task, or subsequent data storage issues. For five additional participants, we excluded their pulse data from our analyses as the pulse-oximeter was unable to acquire a reliable pulse signal. Thus, of the usable runs of functional BOLD data (86 runs across 32 participants), we retained 74.4% of the respiration data (64 runs across 24 participants) and 55.8% of the pulse data (48 runs across 18 participants) for analysis.

We were primarily interested in whether these physiological metrics varied significantly across rewarded and nonrewarded trials in a way that might impact our results. To test this, we calculated trialwise pulse and respiration rates and then compared these rates on rewarded versus nonrewarded trials. After extracting the data for each metric in each run, we used the *peakdet* function in Matlab [84] to locate local minima and maxima in the traces. We then calculated a trialwise rate by counting the number of maxima and minima from the onset of the stimuli on one trial to the onset of the stimuli for the next trial and divided that count by the duration between these two time points (in minutes) to determine the rate. We opted to average the number maxima and minima during the time-window-of-interest in this calculation in order to account for situations where a portion, but not all, of a cardiac or respiratory cycle fell within the window of interest. We then calculated subjectwise averages of these rates for rewarded and nonrewarded trials separately and used a t-test to compare the rates.

### Computational modeling

We conducted an RL modeling analysis on subjects’ trial-by-trial behavioral data. These models allowed us to bridge our behavioral and neural data through the lens of reinforcement learning theory [85]. Specifically, we aimed to model subject behavior to estimate latent variables that reflect the core prediction error processes of RL, and then use these computational estimates to explain variance in neural data using model-based fMRI [86]. This computational analysis thus allowed us to generate inferred reward prediction error (RPE) time-courses for use in our neural analyses.

To model RL behavior and RPEs, we employed a well-established RL model with the basic form:

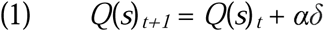

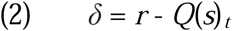

where the value (*Q*) of a given stimulus (*s*) on trial *t* is updated according to the RPE (δ) on that trial (the difference between the expected value *Q*, and received reward *r*), with a learning rate parameter α.

We fit one variant of the model with a single learning rate and one with separate learning rates (α*^+^,* α*^−^*) for rewarded versus nonrewarded outcomes [87,88]. The dual learning rate model was a better fit for participant behavior overall (one learning rate model: summed BIC = 12,182; dual learning rate model: 11,804) so we adopted this model for our analyses:

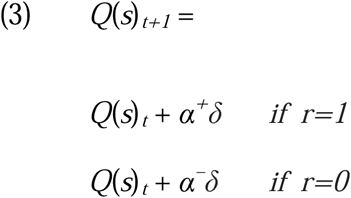

Action selection between the two presented stimuli was modeled using the softmax function:

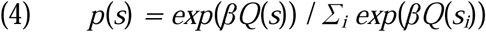

where β reflects the inverse softmax temperature. We used the MATLAB function fmincon to fit our model to each subject’s observed choice data, optimizing the parameter values to maximize the log posterior probability of the choice data given the model. During fitting, α (α*^+^,* α*^−^*) was constrained on [0,1] and β on [0,50], and a Gamma (2,3) prior distribution was used to discourage extreme values of β [88,89].

The fitting procedure was conducted 200 times per subject with randomized starting parameter values to avoid local minima. Simulated choice data (from the optimized model) were produced to investigate and visualize the model’s ability to replicate the main behavioral findings, and to generate the trial-by-trial RPE time courses that were fit to the neural data (see below).

### fMRI acquisition and pre-processing

MR data were acquired with a 3T Siemens Prisma scanner with a 64-channel head coil at BrainWorks at the WuTsai Institute at Yale University. Whole-brain functional images were collected using an echo-planar imaging sequences with Repetition time (TR) = 1000ms, Echo time (TE) = 30ms, voxel size = 2.5×2.5×3 mm^3^, field-of-view = 20.8 x 20.8 x 20.8 cm^3^, 48 slices, P to A phase encoding direction, with multi-band acceleration factor = 3 (interleaved) and in-plane acceleration factor = 2. Gradient echo field maps were acquired to correct for distortions due to B0 inhomogeneities (acquisition parameters: voxel size = 3 × 3 × 3 mm^3^, field-of-view = 24 × 24 × 24 cm^3^). We also collected a high-resolution T1 weighted MPRAGE, voxel size = 1 mm^3^, field-of-view = 25.6 × 25.6 × 25.6 cm^3^. As previously noted, we used Siemens peripherals to track pulse and respiration during the functional runs as well.

### Anatomical data preprocessing

Preprocessing was performed using fMRIPrep 23.2.1 [90,91], which is based on Nipype 1.8.6 [92,93]. The T1w image was corrected for intensity non-uniformity (INU) with N4BiasFieldCorrection [94], distributed with ANTs 2.5.0 [95], and used as T1w-reference throughout the workflow. The T1w-reference was then skull-stripped with a Nipype implementation of the antsBrainExtraction.sh workflow (from ANTs), using OASIS30ANTs as target template. Brain tissue segmentation of cerebrospinal fluid (CSF), white-matter (WM) and gray-matter (GM) was performed on the brain-extracted T1w using fast (FSL, RRID:SCR_002823 [96]). Brain surfaces were reconstructed using recon-all (FreeSurfer 7.3.2, RRID:SCR_001847, [97]), and the brain mask estimated previously was refined with a custom variation of the method to reconcile ANTs-derived and FreeSurfer-derived segmentations of the cortical gray-matter of Mindboggle (RRID:SCR_002438, [98]). Volume-based spatial normalization to one standard space (MNI152NLin2009cAsym) was performed through nonlinear registration with antsRegistration (ANTs 2.5.0), using brain-extracted versions of both T1w reference and the T1w template.

### Functional data preprocessing

For each of the BOLD runs per subject (across all tasks and sessions), the following preprocessing was performed. First, a reference volume was generated, using a custom methodology of fMRIPrep, for use in head motion correction. Head-motion parameters with respect to the BOLD reference (transformation matrices, and six corresponding rotation and translation parameters) are estimated before any spatiotemporal filtering using mcflirt (FSL, [99]). The BOLD reference was then co-registered to the T1w reference using bbregister (FreeSurfer) which implements boundary-based registration [100]. Co-registration was configured with twelve degrees of freedom to account for distortions remaining in the BOLD reference. Several confounding time-series were calculated based on the preprocessed BOLD: framewise displacement (FD), DVARS and three region-wise global signals. FD was computed using two formulations following Power (absolute sum of relative motions, [101]) and Jenkinson (relative root mean square displacement between affines, [99]). FD and DVARS are calculated for each functional run, both using their implementations in Nipype (following the definitions by [101]). The three global signals are extracted within the CSF, the WM, and the whole-brain masks. Additionally, a set of physiological regressors were extracted to allow for component-based noise correction (CompCor, [102]). Principal components are estimated after high-pass filtering the preprocessed BOLD time-series (using a discrete cosine filter with 128s cut-off) for the two CompCor variants: temporal (tCompCor) and anatomical (aCompCor). tCompCor components are then calculated from the top 2% variable voxels within the brain mask. For aCompCor, three probabilistic masks (CSF, WM and combined CSF+WM) are generated in anatomical space. The implementation differs from that of [102] in that instead of eroding the masks by 2 pixels on BOLD space, a mask of pixels that likely contain a volume fraction of GM is subtracted from the aCompCor masks. This mask is obtained by dilating a GM mask extracted from the FreeSurfer’s aseg segmentation, and it ensures components are not extracted from voxels containing a minimal fraction of GM. Finally, these masks are resampled into BOLD space and binarized by thresholding at 0.99 (as in the original implementation). Components are also calculated separately within the WM and CSF masks. For each CompCor decomposition, the k components with the largest singular values are retained, such that the retained components’ time series are sufficient to explain 50 percent of variance across the nuisance mask (CSF, WM, combined, or temporal). The remaining components are dropped from consideration. The head-motion estimates calculated in the correction step were also placed within the corresponding confounds file. The confound time series derived from head motion estimates and global signals were expanded with the inclusion of temporal derivatives and quadratic terms for each [103]. Frames that exceeded a threshold of 0.5 mm FD or 1.5 standardized DVARS were annotated as motion outliers. Additional nuisance time series are calculated by means of principal components analysis of the signal found within a thin band (crown) of voxels around the edge of the brain, as proposed by [104]. All resamplings can be performed with a single interpolation step by composing all the pertinent transformations (i.e. head-motion transform matrices, susceptibility distortion correction when available, and co-registrations to anatomical and output spaces). Gridded (volumetric) resamplings were performed using nitransforms, configured with cubic B-spline interpolation. (Copyright Waiver: Much of the above boilerplate text was automatically generated by fMRIPrep with the express intention that users should copy and paste this text into their manuscripts unchanged. It is released under the CC0 license.) *Additional pre-processing after fMRIprep*

The primary aim of this study was to isolate neural correlates of RL within the cerebellum. Thus, we ran planned analyses on masked cerebellar data, in addition to the whole brain. To constrain analyses within the cerebellum and avoid bleed-over from adjacent visual/temporal regions, we masked the cerebellum from functional data aligned to standard MNI space prior to spatial smoothing. Whole-brain and cerebellar-masked data was spatially smoothed with a 5mm kernel. We excluded runs where participants produced any movement over 2.5mm (1 voxel; N = 10 runs) and excluded participants if they did not have multiple runs to contribute (N = 2). One additional participant was excluded due to technical difficulties. ***fMRI analysis***

### Whole-brain and cerebellar univariate analysis

We modeled BOLD responses using generalized linear models (GLMs) implemented in FSL (version 6.0.7.9). Results from three GLMs are presented in the main text, with additional results from another GLM in the supplement. First, we used a GLM to examine responses to reward versus nonreward feedback. Here, we included separate task regressors, in addition to our confound regressors, for 1) choice stimulus appearance, 2) participant response (or delay onset if there was no response), 3) short-delay feedback (with rewarded trials modeled as 1 and nonrewarded trials as −1), and 4) long-delay feedback (with rewarded trials modeled as 1 and nonrewarded as −1). This design meant that main effects of the feedback regressors would highlight regions of the brain that preferentially respond to rewarding versus nonrewarding feedback. Thus, combinations and contrasts of these two regressors can be used to highlight regions of the brain that are sensitive to reward across delays and also regions that are delay-sensitive.

We used a model-based approach to locate neural correlates of RPEs and surprise (|RPE|) in the brain. To do this, we fit RL models to participant choice data (described above) to obtain participant-specific learning and choice parameters. Then, we used these fitted parameters to simulate trial-by-trial RPEs based on the series of trials that each participant experienced. We included separate parametric regressors for RPEs on short-delay trials versus long-delay trials. In the main version of this model (presented in the main text), we included only rewarded trials in the analysis. Crucially, this allows us to measure true RPE responses that are not confounded with valence. In this model, we additionally included regressors for 1) stimulus appearance, 2) participant response, and 3) the appearance of the feedback. Thus, positive correlations with the RPE regressors should not be due to the appearance of feedback alone or generalized valence responses, but rather with the trial-by-trial variation in inferred RPE. In another version of this model (presented in **Supplemental Figure 3**), we included both rewarded and nonrewarded trials. Here, we included an additional regressor for valence to ensure that correlations with the RPE regressors are not driven by general reward versus nonreward effects (though we note that the valence and RPE regressors are highly colinear, making this model suboptimal). Finally, we took a similar approach to examine correlates of surprise. In this case, the parametric regressor was the absolute value of RPEs. All parametric regressors were z-scored before running the models.

We also conducted a whole-brain GLM with task performance as a covariate in the model to assess if RPE signals were related to behavior. Our primary goal was to assess whether cerebellar RPE activity was related to learning for the short-delay pairs. To that end, we quantified performance for the short-delay pairs by calculating the probability of selecting the higher reward probability shape across all presentations of these pairs. We included this metric as a covariate in the group-level analysis for temporally sensitive RPE signals (i.e., the contrast of RPE activity on rewarded trials on short versus long delay trials). We repeated this analysis with performance for long delay pairs and again for the difference in performance between short and long delay pairs.

In addition to our regressors of interest, we included a variety of motion and noise regressors in all GLMs. We extracted motion regressors from the fMRIprep pipeline and included six rigid body regressors, a DVARS regressor to account for overall motion, and regressors to scrub high motion (>.5mm) TRs. We included the first 10 components of the aCompCor regressors from fMRIprep to account for noise from white matter or cerebrospinal fluid.

Runs were combined within-subject before group-level analyses (FLAME 1). We used a cluster-forming threshold of p < .001 and a family-wise error cluster-corrected threshold of p < 0.001, unless otherwise specified in the text. For cerebellar-specific analyses we cluster-corrected within the cerebellum. We used nilearn (version 0.8.1) to visualize whole-brain results and the SUIT toolbox [105–107] to visualize cerebellar results on flat maps.

### ROI analyses

We conducted additional analyses in *a priori* anatomical ROIs to examine correlates of reward processing and RPE. We used anatomical ROIs from the Harvard-Oxford atlas to create ROI masks. We then extracted subject-wise average /J-parameters from the second-level GLM analysis within each mask for each contrast of interest. We used nonparametric bootstrap hypothesis tests to test whether the measured effect was significantly different from zero in each ROI. We opted for these nonparametric tests as they are robust to statistical assumptions necessary for standard parametric tests. For each bootstrap iteration, we randomly sampled from the measured values with replacement to create a distribution. We repeated this process 1,000 times and computed p-values and 95% confidence intervals by comparing the distribution means to the null value (zero in this case).

### Psychophysiological interaction (PPI) analysis

We used psychophysiological interaction analyses [108] to examine task-dependent changes in functional correlations between the cerebellum and cerebrum during feedback processing. This approach examines whether functional coupling between a seed region and other regions of the brain is modulated by experimental factors. We first extracted a time series of activation from our two *a priori* cerebellar ROIs (Crus I and Crus II) as the physiological regressors in this analysis. Then, we constructed the psychological regressor that identified feedback epochs in the task versus other trial epochs (e.g., stimulus appearance, delay etc.) which was convolved with a canonical hemodynamic response function. Finally, we created a regressor that reflected the interaction of the physiological and psychological regressors. This regressor reveals regions of the brain that significantly covary with the seed region specifically during the epochs of interest.

## Data and code availability statement

Processed data and code will be available upon publication of the manuscript.

## Acknowledgements

We would like to thank the ACT lab and the other Cog Neuro labs at Yale for helpful discussions about this project. These data were collected at BrainWorks at the WuTsai Institute at Yale University. Thank you to Roeland Hancock and Alex Forrence for technical support at BrainWorks and to Tess Levy, Sanghoon Kang, Laurent Caplette, Lily Behm, Diana Wei, Omri Raccah, and Sophie Allen for support with data collection. JET is supported by the NSF GRFP. SDM is supported by NIH grant number R01 NS132926.

