## Supplemental figures for "The human cerebellum encodes temporally sensitive reinforcement learning signals"

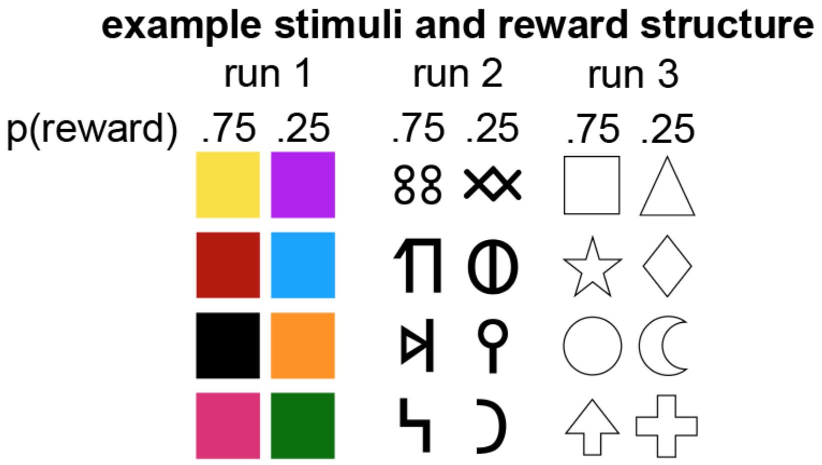


**Supplemental Figure 1.** Example stimulus pairs for three runs of the RL task. One column in each set is associated with a .75 probability of reward and the other with a .25 probability of reward. Importantly stimulus location was randomized across trials such that rewards were not associated with a specific location or action.

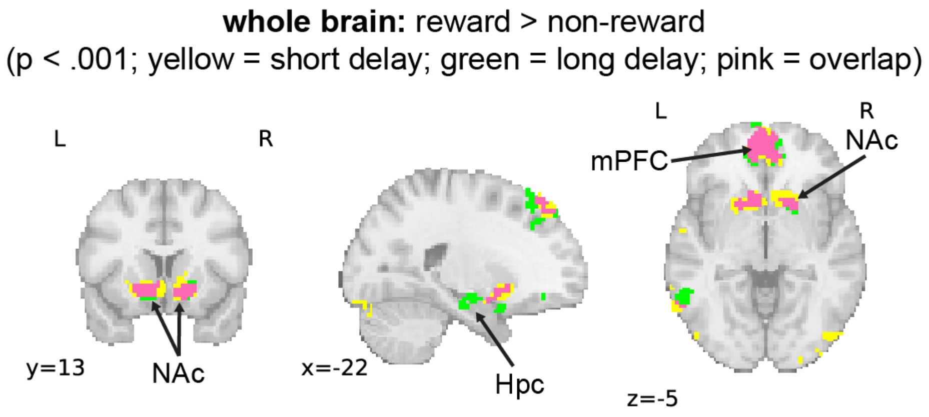


**Supplemental figure 2.** Contrast of activity at the point of feedback for rewarded versus nonrewarded trials in the whole brain (cluster-corrected, p < .001). Yellow represents significant voxels on short delay trials. Green represents significant voxels on long delay trials. Pink voxels are significant on both short and long delay trials.

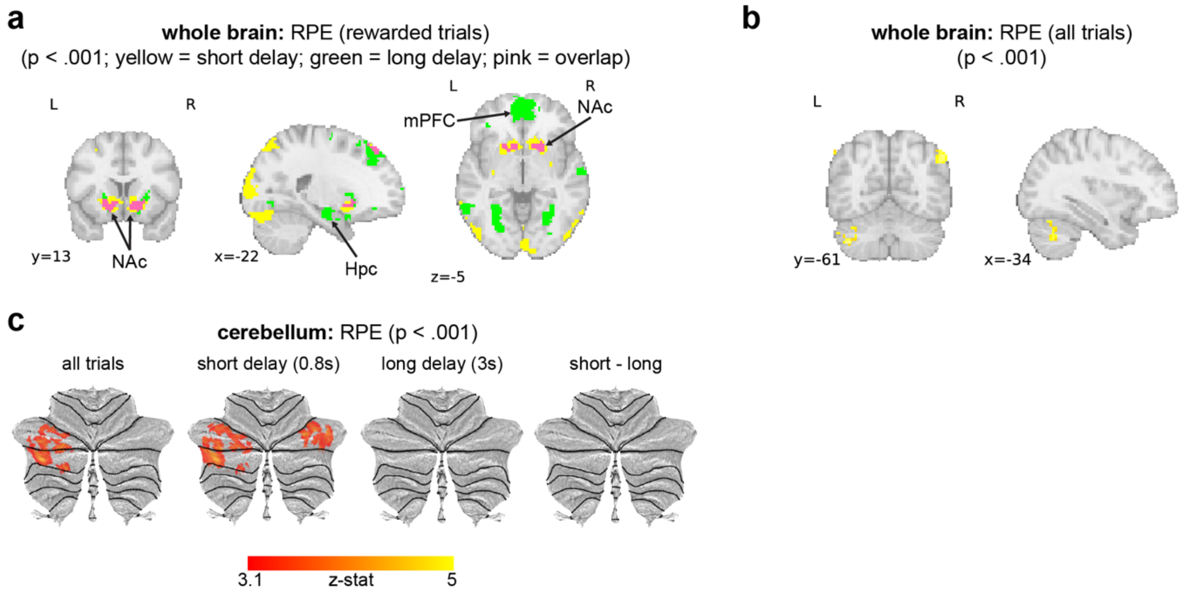


**Supplemental Figure 3.** a) RPE responses on rewarded trials in the whole brain at short and long delays separately (cluster-corrected, p < .001). Yellow represents significant voxels on short delay trials. Green represents significant voxels on long delay trials. Pink voxels were significant on both short and long delay trials. b) Significant RPE results in the whole brain across rewarded and nonrewarded trials. c) RPE responses in cerebellum across rewarded and nonrewarded trials. First panel combines across delay intervals and subsequent panels plot responses at short and long delays separately and their interaction.
